# Disrupted basal ganglia output during movement preparation in hemi-parkinsonian mice accounts for behavioral deficits

**DOI:** 10.1101/2020.06.19.160457

**Authors:** Anand Tekriwal, Mario J. Lintz, John A. Thompson, Gidon Felsen

**Author notes:** To whom correspondence should be addressed, Department of Physiology and Biophysics, University of Colorado School of Medicine, 12800 E. 19th Ave., Mail Stop 8307, Aurora, CO 80045, USA.

## Abstract

Parkinsonian motor deficits are associated with elevated inhibitory output from the basal ganglia (BG). However, several features of Parkinson’s disease (PD) have not been accounted for by this simple “rate model” framework, including the observation in PD patients that movements guided by external stimuli are less impaired than otherwise-identical movements generated based on internal goals. Is this difference in impairment due to divergent processing within the BG itself, or to the recruitment of extra-BG pathways by sensory processing? In addition, surprisingly little is known about precisely when, in the sequence from selecting to executing movements, BG output is altered by PD. Here, we address these questions by recording activity in the SNr, a key BG output nucleus, in hemiparkinsonian (hemi-PD) mice performing a well-controlled behavioral task requiring stimulus-guided and internally-specified directional movements. We found that hemi-PD mice exhibited a bias ipsilateral to the side of dopaminergic cell loss that was stronger when movements were internally specified rather than stimulus guided, consistent with clinical observations in parkinsonian patients. We further found that changes in parkinsonian SNr activity during movement preparation could account for the ipsilateral behavioral bias, as well as its greater magnitude for internally-specified movements, consistent with some aspects of the rate model. These results suggest that parkinsonian changes in BG output underlying movement preparation contribute to the greater deficit in internally-specified than stimulus-guided movements.

## Introduction

Parkinson’s disease (PD) is a neurodegenerative disease of the basal ganglia (BG) in which motor impairments arise from disordered – typically, elevated – inhibitory BG output resulting from the loss of dopaminergic tone (DeLong, 1990; Wichmann et al., 1999; Ibanez-Sandoval et al., 2007; Utter and Basso, 2008; Wang et al., 2010a; Seeger-Armbruster and von Ameln-Mayerhofer, 2013; Brazhnik et al., 2014; Filyushkina et al., 2019; McGregor and Nelson, 2019). One predominant theoretical framework for BG pathology in PD is the “rate model”, which posits that motor centers downstream of the BG are over-inhibited, leading to disordered movements (Albin et al., 1989; DeLong, 1990; Obeso et al., 2008; Utter and Basso, 2008; McGregor and Nelson, 2019; Vitek and Johnson, 2019). However, it is not clear whether the rate model can account for context-dependent PD motor phenomena, including the intriguing clinical observation that not all forms of movement are equally affected by PD: when movements are guided by external stimuli (e.g., gait matching with a rhythmic auditory stimulus or visually patterned flooring, kinematics are less impaired than for otherwise identical movements made in the absence of guiding stimuli (Glickstein and Stein, 1991; McIntosh et al., 1997; Ballanger et al., 2006; Daroff, 2008; McDonald et al., 2015; Distler et al., 2016). The primary question raised by this observation is whether parkinsonian BG output is similarly disrupted for these “stimulus-guided” and “internally-specified” movements. If so, we might infer that stimulus-guided movements are protected from PD via the recruitment of extra-BG pathways (Lewis et al., 2007; Hackney et al., 2015; Drucker et al., 2019; Filyushkina et al., 2019; Chen et al., 2020). However, differences in parkinsonian BG output between these forms of movements – particularly differences consistent with rate model predictions – would implicate BG processing itself in this behavioral phenomenon. Given that understanding the neural basis for this clinical observation could be leveraged to improve treatment for PD, we sought to develop an experimental paradigm for examining parkinsonian BG output during stimulus-guided and internally-specified movements.

We focused on parkinsonian BG output during movement *preparation*, a key motor phase in which sensory and cognitive variables are integrated (Cisek and Kalaska, 2010), and when disrupted may contribute to bradykinesia (Dick et al., 1984; Jahanshahi et al., 1992; Suri et al., 1998; Berardelli et al., 2001; Cutsuridis and Perantonis, 2006; Moroney et al., 2008; Wu et al., 2015; Hess and Hallett, 2017). Under normal conditions, the substantia nigra pars reticulata (SNr), a BG output nucleus, is strongly engaged by the preparation of directional movements (Handel and Glimcher, 1999; Sato and Hikosaka, 2002; Lintz and Felsen, 2016). However, while numerous studies have examined parkinsonian changes in SNr activity under passive conditions or rhythmic locomotion (Hutchison et al., 1994; Wichmann et al., 1999; Galati et al., 2010; Wang et al., 2010a; Seeger-Armbruster and von Ameln-Mayerhofer, 2013; Brazhnik et al., 2014; Lobb and Jaeger, 2015; Aristieta et al., 2016; Willard et al., 2019), this approach is insufficient for differentiating how SNr activity during distinct motor phases, including movement preparation, is affected by PD. To address this question, we recorded SNr activity during a behavioral task in which mice with unilateral dopaminergic cell loss prepare, and subsequently initiate, SNr-engaging directional (left or right) movements that are either stimulus-guided or internally-specified (Uchida and Mainen, 2003; Thompson and Felsen, 2013; Lintz and Felsen, 2016). Crucially, by requiring that mice wait for a go signal before initiating their movement, movement preparation is temporally isolated from initiation, allowing the dissociation of PD impact on BG output underlying these processes.

We found that mice exhibited a directional bias ipsilateral to the hemisphere with dopaminergic cell loss that was more prominent on internally-specified than stimulus-guided trials, accordant with clinical observations of context-dependent motor effects in PD. Furthermore, we found that SNr activity during movement preparation was altered in a manner consistent with some, but not all, rate-model predictions about the relationship between BG output and behavior, suggesting that reorganization of BG processing by dopaminergic cell loss contributes to the greater deficit in performance of internally-specified than stimulus-guided movements. These findings inform our understanding of BG pathophysiology and can contribute to refining neuromodulatory PD treatments.

## Materials and Methods

### Animal subjects

All experiments were performed according to protocols approved by the University of Colorado Anschutz Medical Campus Institutional Animal Care and Use Committee. Subjects were male adult C57BL/6J mice (aged 7–14 months at the start of experiments; Jackson Labs) housed in a vivarium with a 12-hr light/dark cycle with lights on at 5:00 am. Food (Teklad Global Rodent Diet No. 2918; Harlan) was available *ad libitum*. Access to water was restricted prior to the behavioral session to motivate performance; however, if mice did not obtain ~1 ml of water during the behavioral session, additional water was provided for ~2–5 min following the behavioral session. All mice were weighed daily and received sufficient water during behavioral sessions to maintain >85% of pre-water restriction weight.

For behavioral analyses, only mice that completed at least 15 pre- and 15 post-surgery sessions were included (hemiparkinsonian (hemi-PD), n = 4; control, n = 4). For electrophysiological analyses, mice were included if well-isolated neurons were recorded during the task (hemi-PD, n = 5; control, n = 4). Some data from control mice were previously published using different analyses than the current study (Lintz and Felsen, 2016). For rotation assay analyses, only mice that completed at least three pre- and three-post surgery rotation assay sessions were included (hemi-PD, n = 5; control, n = 3).

### Behavioral task

Mice were trained on a task requiring stimulus-guided (SG) and internally-specified (IS) movements (Fig. 1) as previously described (Lintz and Felsen, 2016). Briefly, each mouse was water-restricted and trained to interact with three ports (center: odor port; sides: reward ports; Fig. 2A) along one wall of a behavioral chamber (Island Motion). All behavior was performed in the dark in the absence of visual cues. On each trial, the mouse entered the odor port, triggering the delivery of an odor; waited 488 ± 104 ms (mean ± SD) for an auditory go signal; exited the odor port; and entered one of the reward ports (Fig. 2A). Premature exit from the odor port resulted in the unavailability of reward on that trial. Odors were comprised of binary mixtures of (-)-carvone (“Odor A”) and (+)-carvone (“Odor B”). On each SG trial, one of seven odor mixtures was presented via an olfactometer (Island Motion): Odor A/Odor B = 95/5, 80/20, 60/40, 50/50, 40/60, 20/80, or 5/95. Mixtures in which Odor A > Odor B indicated reward availability only at the left port, mixtures in which Odor B > Odor A indicated reward availability only at the right port and for mixtures in which Odor A = Odor B (i.e., the 50/50 mixture) reward was equally likely (probability = 0.5) at both ports (Fig. 2B). Since we surgically targeted the left hemisphere in all mice, we refer to Odor A as the “ipsilateral odor” and Odor B as the “contralateral odor” (e.g., Fig. 2C). Similarly, we refer to the directions “left” and “right” as “ipsilateral” and “contralateral”, respectively. On trials in which Odor A = Odor B (Odor A/Odor B = 50/50), the probability of reward at the ipsilateral and contralateral ports, independently, was 0.5. Reward, consisting of 4 μl of water, was delivered by transiently opening a calibrated water valve 10–100 ms after reward port entry. Odor and water delivery were controlled, and port entries and exits were recorded, using custom software (available at https://github.com/felsenlab; adapted from C. D. Brody) written in MATLAB (MathWorks).

**Figure 1.**
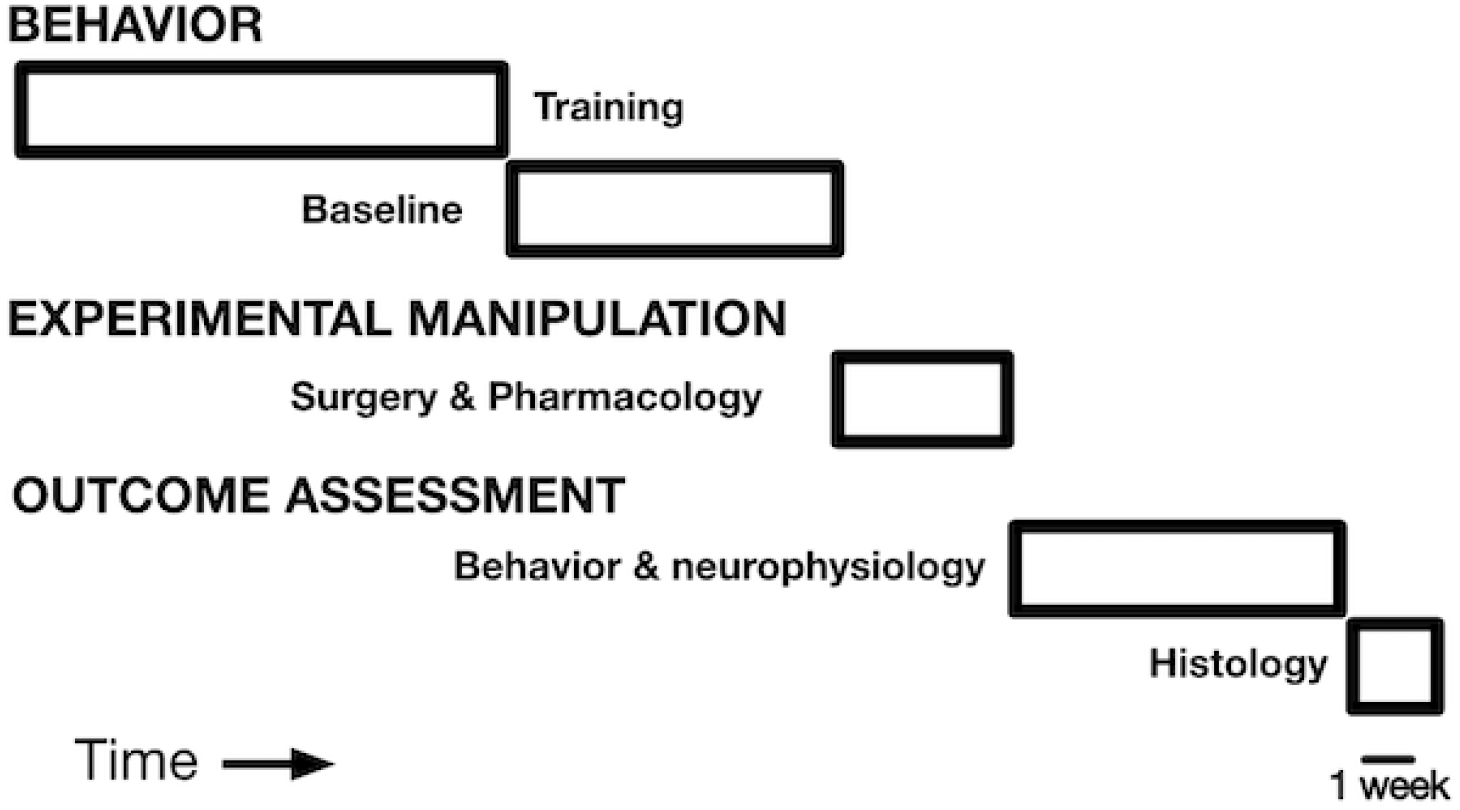
Experimental timeline.

**Figure 2.**
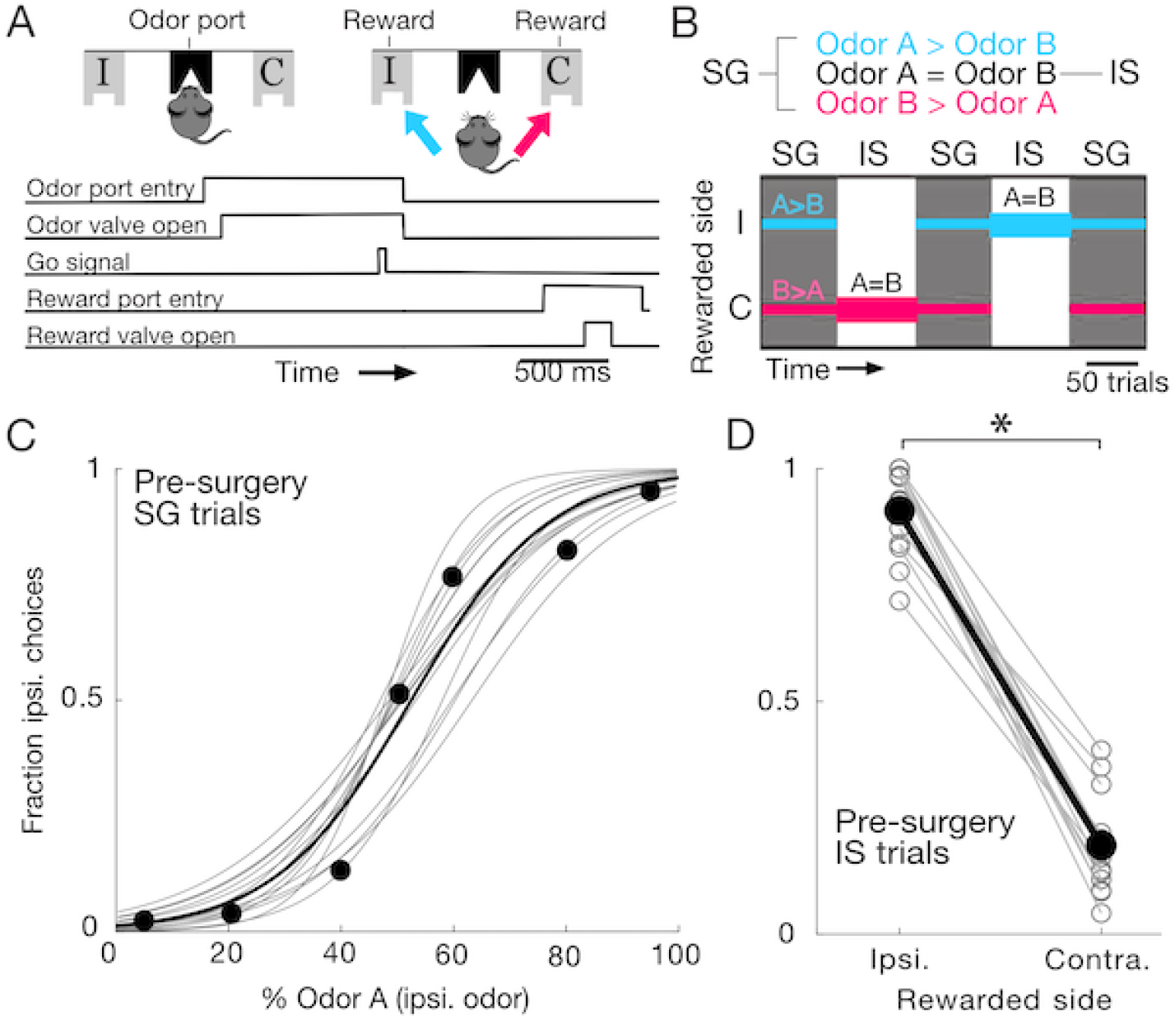
Behavioral task and baseline performance. ***A*,** Port locations (top) and timing of task events (bottom). Ipsilateral (I) and contralateral (C) are defined relative to the side of brain targeted for surgery (always left). Cyan represents ipsilateral choices, magenta represents contralateral choices. ***B***, Odor mixtures on SG (stimulus-guided) and IS (internally-specified) trials (top) and interleaving of SG and IS blocks within a session (bottom). On SG trials, the Odor A/Odor B mixture presented was 5/95, 20/80, 40/60, 50/50, 60/40, 80/20 or 95/5. On IS trials, the Odor A/Odor B mixture presented was always 50/50. Horizontal cyan and magenta lines indicate which port(s) were rewarded in each block: On SG trials, ipsilateral side was rewarded when Odor A > Odor B (cyan), contralateral side was rewarded when Odor B > Odor A (magenta), and either side was equally likely to be rewarded when Odor A = Odor B; on all IS trials, Odor A = Odor B and only one side was rewarded throughout the block. ***C*,** Baseline (pre-surgery) performance on SG trials for representative mouse subsequently assigned to the hemi-PD group. Gray lines show best-fit logistic functions (p = 1/1 + *e*^(−*a* −*bx*)^ where *x* is the proportion of Odor A, *p* is the fraction of ipsilateral choices, and *a* and *b* are the best-fit free parameters) for each session (n = 17). Circles show average across sessions for the binary odor mixtures presented; solid line shows best-fit logistic function to all choices across sessions. ***D***, Baseline (pre-surgery) performance on IS trials for representative mouse subsequently assigned to hemi-PD group. Gray lines link ipsilateral- and contralateral-rewarded IS blocks within the same session (n = 15). Black circles indicate medians. Mouse chose the ipsilateral port more often on ipsilateral-rewarded blocks (p = 3.05 × 10^-5^, 1-tailed Wilcoxon signed rank test).

Mice learned to perform SG trials (Fig. 2C) within ~48 sessions (1 session/day); detailed training stages are described in Stubblefield et al. (Stubblefield et al., 2013). Mice required an additional ~5 sessions to learn to perform interleaved blocks of SG and IS trials. On each IS trial the 50/50 mixture was presented, and reward was available only at one side throughout the block (Fig. 2D). Detailed training stages for IS trials are described in Lintz & Felsen (Lintz and Felsen, 2016). Mice performed 5 blocks (SG, IS, SG, IS, SG) per session (Fig. 2B); the side associated with reward switched between each IS block. Upon completing training, mice performed at least 15 sessions to establish pre-surgery baseline behavior, underwent surgery (see below), and subsequently resumed task performance, beginning with sessions consisting of SG trials only (post-surgery behavior; Fig. 1).

### Rotation assay

The direction of spontaneous movement was assessed before and after surgery using a standard rotation assay (Ungerstedt, 1976; Smith and Heuer, 2011). Following intraperitoneal (i.p.) administration of d-amphetamine (2.5 mg/kg, Sigma), mice were placed in a transparent beaker with a diameter of 11.5 cm. Mice were monitored for the next 90 minutes and behavior recorded using an overhead camera. Rotations were analyzed from 10 to 30 minutes post i.p. injection. A rotation score was calculated by counting the total number of complete ipsilateral (left) rotations and subtracting the total number of complete contralateral (right) rotations. Repeated testing was carried out with at least 1 week between d-amphetamine injections to allow for recovery.

### Substantia nigra pars compacta surgery – Unilateral 6 OHDA and saline injections

The mouse was anesthetized with isoflurane and secured in a stereotaxic device, the skull was exposed with a midline incision, and a craniotomy targeting the left SNc (substantia nigra pars compacta) was performed, centered at 3.07 mm posterior from bregma, 1.250 mm lateral from the midline, and 4.35 mm deep from cortical surface (Paxinos and Franklin, 2004). Injection volume totaled 2 μL injected at target (2.43 mg 6-OHDA/mL 0.02% ascorbic acid). After suturing the incision, a topical triple antibiotic ointment (Major) mixed with 2% lidocaine hydrochloride jelly (Akorn) was applied to the scalp, the mouse was removed from the stereotaxic device, the isoflurane was turned off, and oxygen alone was delivered to the mouse to gradually alleviate anesthetic state. Mice were administered sterile isotonic saline (0.9%) for rehydration and an analgesic (Ketofen; 5 mg/kg) for pain management. Analgesic and topical antibiotic administration was repeated daily for up to 5 days, and mice were closely monitored for any signs of distress.

This procedure was identical for control mice assessed on the rotation assay but with saline injected instead of 6-OHDA. These mice did not undergo further surgery and were used solely for rotation assay testing.

### SNr surgery – tetrode implantation

Details of the surgical procedure are provided in Thompson and Felsen (Thompson and Felsen, 2013). Briefly, following establishment of pre-surgery baseline behavior and (in hemi-PD mice) following unilateral 6-OHDA injections (Fig. 1), the mouse was anesthetized with isoflurane and secured in a stereotaxic device, the scalp was incised and retracted, 2 small screws were attached to the skull, and a craniotomy targeting the left SNr was performed, centered at 3.07 mm posterior from bregma and 1.25 mm lateral from the midline (Franklin and Paxinos, 2004). A VersaDrive 4 microdrive (Neuralynx), containing 4 independently adjustable tetrodes, was affixed to the skull via the screws, luting (3M), and dental acrylic (A-M Systems). A second small craniotomy was performed in order to place the ground wire in direct contact with the brain. After the acrylic hardened, a topical triple antibiotic ointment (Major) mixed with 2% lidocaine hydrochloride jelly (Akorn) was applied to the scalp, the mouse was removed from the stereotaxic device, the isoflurane was turned off, and oxygen alone was delivered to the mouse to gradually alleviate anesthetic state. Mice were administered sterile isotonic saline (0.9%) for rehydration and an analgesic (Ketofen; 5 mg/kg) for pain management. Analgesic and topical antibiotic administration was repeated daily for up to 5 days, and mice were closely monitored for any signs of distress.

### Electrophysiology

Neural recordings were collected using four tetrodes, wherein each tetrode consisted of four polyimide-coated nichrome wires (Sandvik; single-wire diameter 12.5፧μm) gold plated to 0.2–0.4 MΩ impedance. Electrical signals were amplified and recorded using the Digital Lynx S multichannel acquisition system (Neuralynx) in conjunction with Cheetah data acquisition software (Neuralynx). Tetrode depths were adjusted approximately 23 hours before each recording session in order to sample an independent population of neurons across sessions. To estimate tetrode depths during each session we calculated distance traveled with respect to the rotation fraction of the screw that was affixed to the shuttle holding the tetrode. One full rotation moved the tetrode ~250፧μm and tetrodes were moved ~62.5፧μm between sessions. The final tetrode location was confirmed through histological assessment (see below).

Offline spike sorting and cluster quality analysis was performed using MClust software (MClust-4.3, A.D. Redish, et al.) in MATLAB. Briefly, for each tetrode, single units were isolated by manual cluster identification based on spike features derived from sampled waveforms (see example mean waveforms recorded in control mice in Lintz & Felsen, 2016). Identification of single units through examination of spikes in high-dimensional feature space allowed us to refine the delimitation of identified clusters by examining all possible two-dimensional combinations of selected spike features. We used standard spike features for single unit extraction: peak amplitude, energy (square root of the sum of squares of each point in the waveform, divided by the number of samples in the waveform), and the first principal component normalized by energy. Spike features were derived separately for individual leads. To assess the quality of identified clusters we calculated two standard quantitative metrics: L-ratio and isolation distance (Schmitzer-Torbert et al., 2005). Clusters with an L-ratio of less than 0.75 and isolation distance greater than 6.5 were deemed single units, which resulted in the exclusion of 7% of the identified clusters. Only clusters with few interspike intervals less than 1.5 ms were considered for further examination. Furthermore, we excluded the possibility of including data from the same neuron twice by ensuring that both the waveforms and response properties sufficiently changed across sessions. If they did not, we conservatively assumed that we were recording from the same neuron, and only included data from one session.

### Immunohistochemistry

Final tetrode locations were verified by producing electrolytic lesions (100 mA, −1.5 min per lead) after the last recording session (Lintz and Felsen, 2016). Mice were then overdosed with an i.p. injection of sodium pentobarbital (100 mg/kg) and transcardially perfused with saline followed by ice-cold 4% paraformaldehyde (PFA) in 0.1፧M phosphate buffer (PB). After perfusion, brains were submerged in 4% PFA in 0.1፧M PB for 24 hours for post-fixation and then cryoprotected for 24 hours by immersion in 30% sucrose in 0.1፧M PB. The brain was encased in the same sucrose solution, and frozen rapidly on dry ice.

Serial coronal sections (60፧μm) were cut on a sliding microtome. Fluorescent Nissl (NeuroTrace, Invitrogen) was used to identify cytoarchitectural features of the SNr and verify tetrode tracks and lesion damage within or below the SNr, as previously described (Lintz and Felsen, 2016). In addition, coronal sections were stained for tyrosine hydroxylase (TH). Following repeated soaks in PBS and blocking solution, sections were exposed to primary antibody overnight (Anti-Tyrosine Hydoxylase (Rabbit) Antibody, 1:1000, Rockland). Next, sections were washed in carrier solution (2×10-min) and exposed to secondary antibody for 2 hours (Goat anti-Rabbit IgG (H+L) Secondary Antibody, 1:500). Images were captured with a 10x objective lens, using an LSM 5 Pascal series Axioskop 2 FS MOT confocal microscope (Zeiss). For each mouse, a representative coronal section including the SNc (Paxinos & Franklin) was used to quantify dopaminergic cell loss by comparing the number of TH+ neurons ipsilateral and contralateral to the injection.

Dopaminergic cell loss was quantified by calculating the percent decrease of red (TH+) pixel intensity in the SNc on the injected side relative to the SNc on the non-injected side, after accounting for background differences in red pixel intensity between the two sides. Hemi-PD mice without verified >70% dopaminergic cell loss (5/10), were excluded from the group. Of the excluded mice, 4 were due to missing or insufficient tissue without which TH+ could not be confirmed while the remaining mouse was excluded for <70% loss (65%). Average TH+ loss of the 5 hemi-PD mice ranged from 73% to 88%, with a median of 78%. Secondary confirmation of dopaminergic cell loss was quantified in the same manner using coronal sections containing the striatum (Fig. 3A).

**Figure 3.**
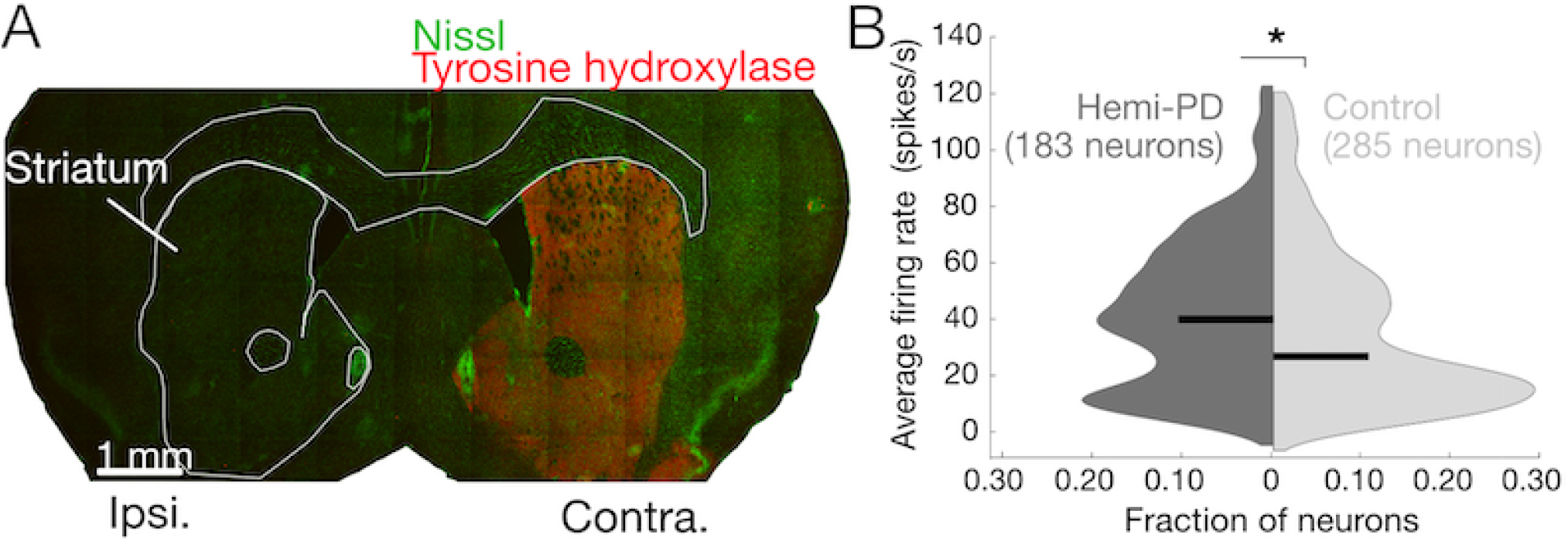
Validation of hemi-PD mouse model. ***A***, Representative coronal section (0.97 mm anterior to Bregma) in hemi-PD mouse. 6-OHDA was delivered to the ipsilateral SNc. Green, Nissl; red, tyrosine hydroxylase. ***B***, Baseline activity (odor port entry to reward port exit in each trial) of SNr neurons during task performance in hemi-PD (dark gray, n = 5) and control (light gray, n = 4) mice. Black lines, medians. Median baseline SNr activity was higher in hemi-PD mice (p = 0.0157, 1-tailed Wilcoxon rank sum test).

### Behavioral directional bias

Behavioral directional bias was quantified as the difference between the fraction of ipsilateral choices on ipsilaterally-rewarded trials (i.e., trials for which reward was available at the ipsilateral port) and the fraction of contralateral choices on contralaterally-rewarded trials (i.e., trials for which reward was available at the contralateral port):

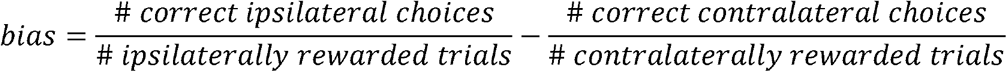

Positive values reflect an ipsilateral behavioral bias and negative values reflect a contralateral bias. 50/50 SG trials were evenly split between ipsilaterally-rewarded and contralaterally-rewarded and either choice was considered correct. To compare directional bias between presurgery and post-surgery sessions (Figs. 4, 5), we calculated a single pre-surgery bias for each mouse based on all pre-surgery trials performed. To compare directional bias between SG and IS trials within the same session (Fig. 6), to account for ceiling effects on ipsilaterally-rewarded IS trials, the bias calculation was modified to quantify ipsilateral choices on contralaterally-rewarded trials only. Sessions were included in this analysis if the mouse completed at least 25 trials of each direction (ipsilateral and contralateral) and type (SG and IS) trials.

**Figure 4.**
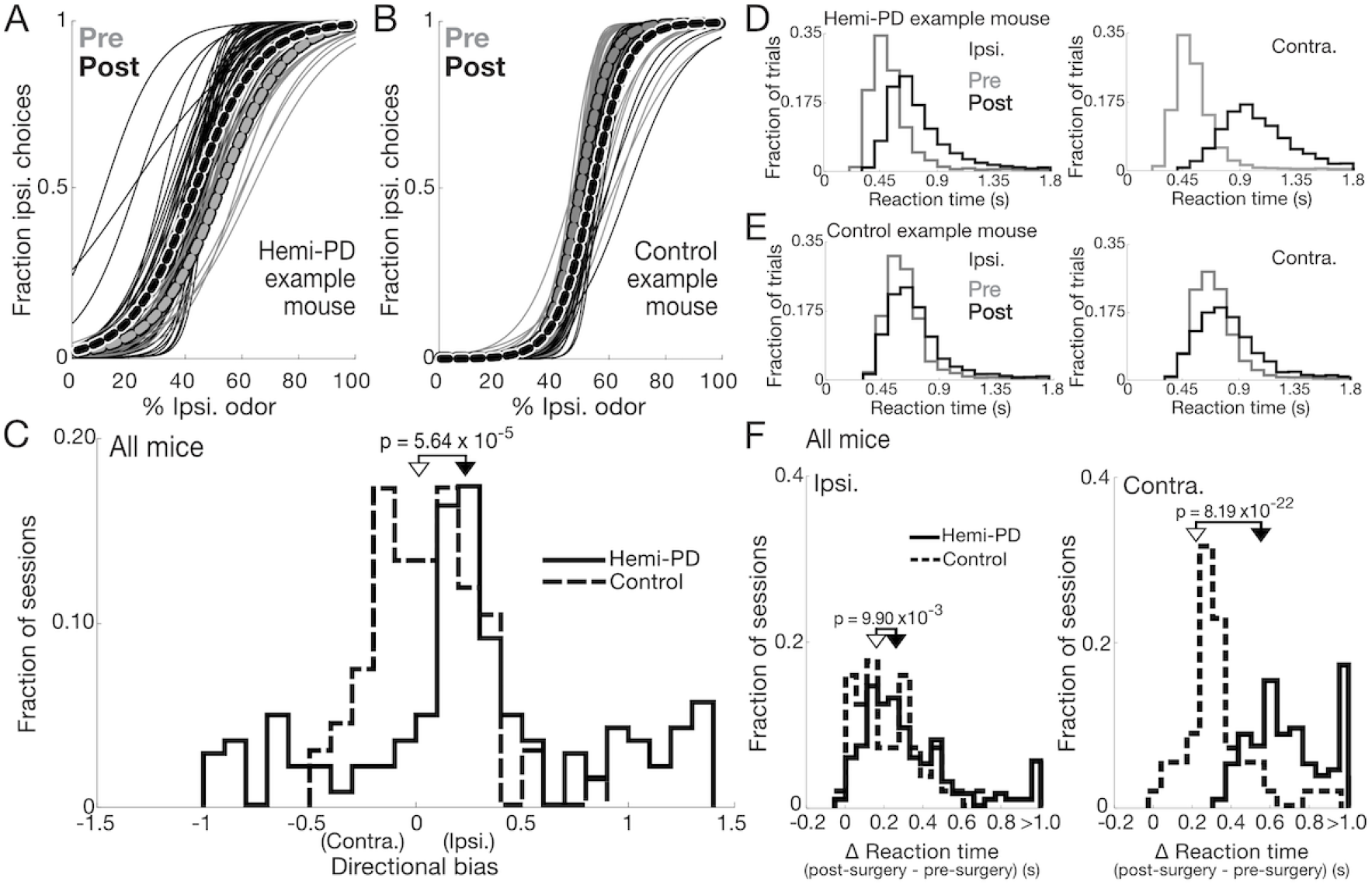
Effect of unilateral dopaminergic cell loss on stimulus-guided (SG) movements. ***A*,** Thin lines show best-fit logistic functions (as in Fig. 2C) for SG trials for each pre-surgery session (gray, n = 30) and post-surgery session (black, n = 42) for a representative hemi-PD mouse. Thick lines show fits for SG trials combined across all pre-surgery (grey) and post-surgery (black) sessions, ***B*,** As in A, for a representative control mouse (tetrode drive implanted to SNr; n = 33 pre-surgery and 17 post-surgery sessions). ***C*,** Directional bias (relative to the pre-surgery baseline for each mouse) on SG trials in each post-surgery session (138 sessions in 4 hemi-PD mice; 57 sessions in 4 control mice). Hemi-PD mice exhibited an ipsilateral bias (p = 8.87 × 10^-8^, 2-tailed Wilcoxon signed rank test); control mice did not (p = 0.3480, 2-tailed t-test; post-surgery bias differed between hemi-PD than control mice, p = 5.64 × 10^-5^, 2-tailed Wilcoxon rank sum test; black and white arrow indicates median values for hemi-PD and control mice, respectively). ***D*,** Reaction times (from go signal to reward port entry) on ipsilateral (left panel) and contralateral (right panel) SG trials in pre-surgery (gray) and post-surgery (black) trials for a representative hemi-PD mouse (ipsilateral: 1,957 trials pre-surgery, 5,393 trials post-surgery; contralateral: 2,193 trials pre-surgery, 3,143 trials post-surgery). ***E*,** As in ***D*,** for a representative control mouse (ipsilateral: 1,386 trials pre-surgery, 2,340 trials post-surgery; contralateral: 1,621 trials pre-surgery, 1,760 trials post-surgery). ***F*,** Change (from the pre-surgery median baseline for each mouse) in median reaction time on each post-surgery session (same mice and sessions as in C) on ipsilateral (left panel) and contralateral (right panel) SG trials. Hemi-PD mice exhibited larger changes in reaction times than control mice (ipsilateral trials: p = 9.90 × 10^-3^, contralateral trials: p = 8.19 × 10^-22^, 2-tailed Wilcoxon rank sum tests; black and white arrow indicates median values for hemi-PD and control mice, respectively), and exhibited a larger change on contralateral than on ipsilateral trials (p = 1.66 × 10^-12^, 2-tailed Wilcoxon signed rank test).

**Figure 5.**
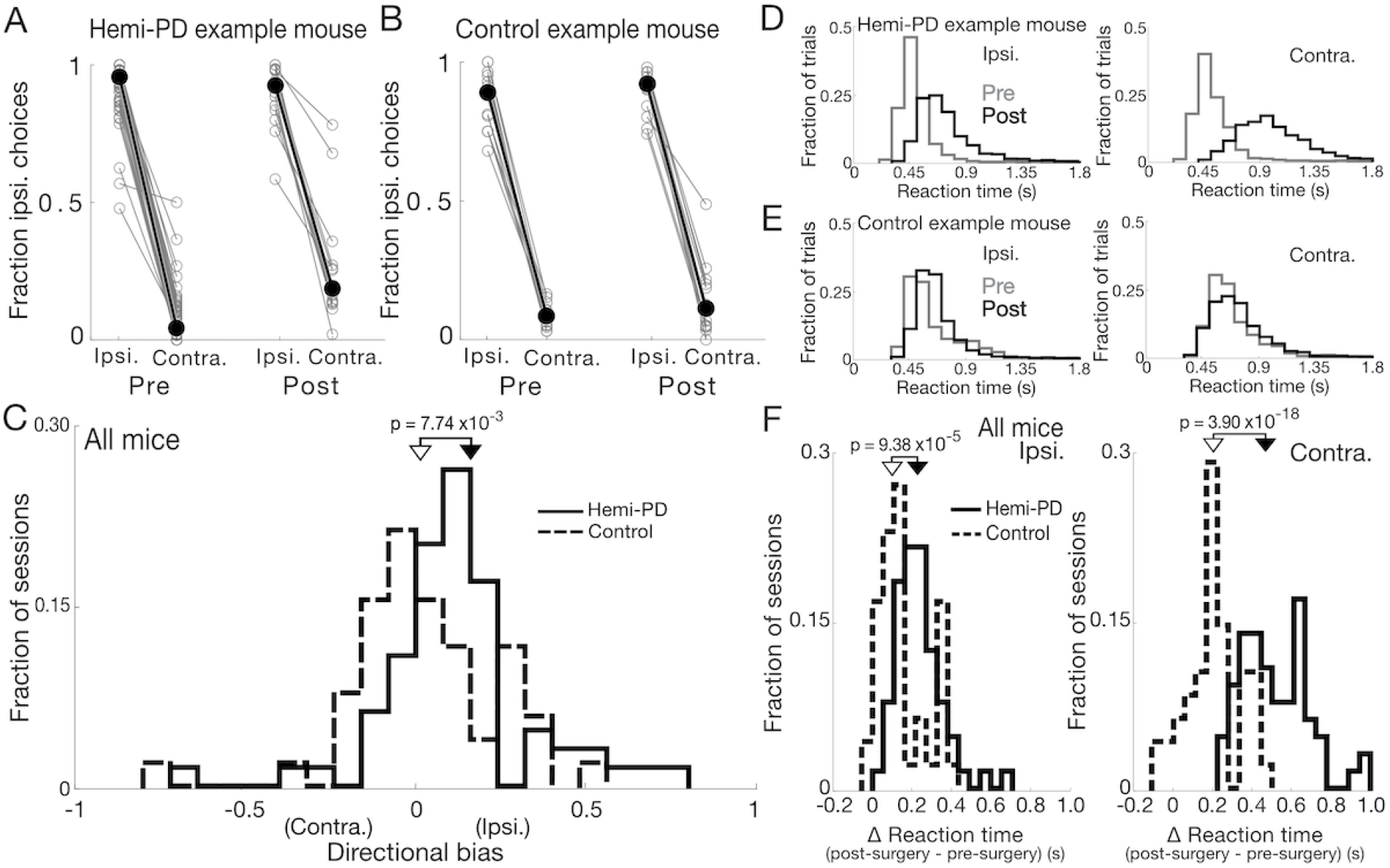
Effect of unilateral dopaminergic cell loss on internally-specified (IS) movements. ***A***, Connected symbols show fraction of ipsilateral choices on ipsilateral and contralateral blocks of IS trials for each pre-surgery session (n = 63 sessions) and post-surgery session (n = 16) for a representative hemi-PD mouse. Filled symbols show means. ***B***, As in ***A***, for a representative control mouse (tetrode drive implanted; n = 14 pre-surgery and 14 post-surgery sessions). ***C*,** Directional bias (relative to the pre-surgery baseline for each mouse) on IS trials in each post-surgery session (65 sessions in 4 hemi-PD mice; 48 sessions in 4 control mice). Hemi-PD mice exhibited an ipsilateral bias (p = 1.49 × 10^-6^, 2-tailed Wilcoxon signed rank test); control mice did not (p = 0.3855, 2-tailed t-test; post-surgery bias differed between hemi-PD and control mice, p = 7.74 × 10^-3^, 2-tailed Wilcoxon rank sum test; black and white arrow indicates median values for hemi-PD and control mice, respectively). ***D*,** Reaction times (from go signal to reward port entry) on ipsilateral (left panel) and contralateral (right panel) IS trials in pre-surgery (gray) and post-surgery (black) trials for a representative hemi-PD mouse (ipsilateral: 874 trials pre-surgery, 2,328 trials post-surgery; contralateral: 789 trials pre-surgery, 1,783 trials post-surgery). ***E*,** As in ***D*,** for a representative control mouse (ipsilateral: 764 trials pre-surgery, 839 trials post-surgery; contralateral: 820 trials pre-surgery, 804 trials post-surgery). ***F*,** Change (from the pre-surgery median baseline for each mouse) in median reaction time on each post-surgery session (same mice and sessions as in C) on ipsilateral (left panel) and contralateral (right panel) IS trials. Hemi-PD mice exhibited larger changes in reaction times than control mice (ipsilateral trials: p = 9.38 × 10^-5^, 2-tailed Wilcoxon rank sum test, contralateral trials: p = 3.90 × 10^-18^, 2-tailed t-test; black and white arrow indicates median values for hemi-PD and control mice, respectively), and exhibited a larger change on contralateral than on ipsilateral trials (p = 1.09 × 10^-10^, 2-tailed Wilcoxon signed rank test).

**Figure 6.**
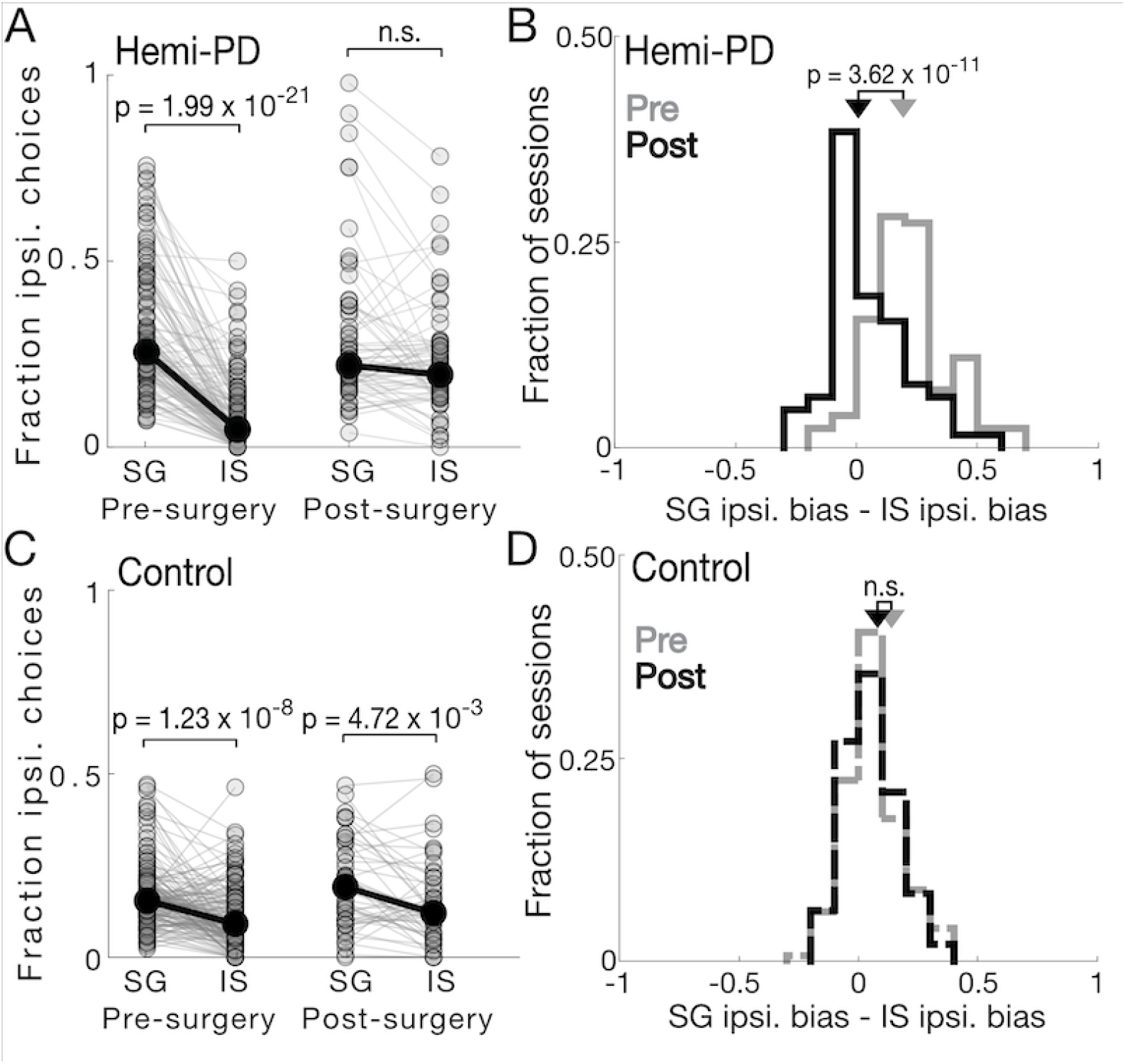
Direct comparison between stimulus-guided (SG) and internally-specified (IS) behavior in hemi-PD mice. ***A*,** Connected symbols show fraction of ipsilateral choices on SG and IS contralaterally-rewarded trials within each pre-surgery session (n = 128) and post-surgery session (n = 65) for hemi-PD mice (n = 4). Hemi-PD mice were more ipsilaterally biased on IS than SG trials pre-surgery (p = 1.99 × 10^-21^, 2-tailed Wilcoxon signed rank test), but not post-surgery (p = 0.0924, 2-tailed Wilcoxon signed rank test). ***B***, Within-session ipsilateral bias differences between SG and IS trials changed between pre- and post-surgery sessions (p = 3.62 × 10^-11^, 2-tailed Wilcoxon rank sum test). C, As in A, for control mice (n = 148 pre-surgery sessions, n = 48 post-surgery sessions, n = 4 mice). Control mice were more ipsilaterally biased on IS than SG trials pre-surgery (p = 1.23 × 10^-8^, 2-tailed Wilcoxon signed rank test) and postsurgery (p = 4.72 × 10^-3^, 2-tailed Wilcoxon signed rank test). ***D*,** As in B, for control mice. Within-session ipsilateral bias differences between SG and IS trials did not change between pre- and post-surgery sessions (p = 0.643, 2-tailed Wilcoxon rank sum test).

### Neuronal direction preference

We used a firing-rate-based ROC-based analysis to quantify the selectivity of single neurons for movement direction (Green and Swets, 1966; Lintz and Felsen, 2016). This analysis calculates the ability of an ideal observer to classify whether a given firing rate was recorded in one of two conditions (i. e., preceding ipsilateral (left) or contralateral (right) movement). We defined “preference” as 2(ROC_area_ – 0.5), a measure ranging from −1 to 1, where −1 signifies the strongest possible preference for ipsilateral, 1 signifies the strongest possible preference for contralateral, and 0 signifies no preference (Feierstein et al., 2006; Lintz and Felsen, 2016). For example, if the firing rate of a given neuron is generally higher preceding ipsilateral than contralateral movements, that neuron is assigned a preference < 0. Statistical significance was determined with a permutation test: we recalculated the preference after randomly reassigning all firing rates to either of the two groups, repeating this procedure 500 times to obtain a distribution of values, and calculated the fraction of random values exceeding the actual value. We tested for significance at α = 0.05. Trials in which the movement time (between odor port exit and reward port entry) was > 1.5፧S were excluded from all analyses. Neurons with < 25 trials of each trial type under comparison (ipsilateral SG, contralateral SG, ipsilateral IS, or contralateral IS) or with a firing rate < 2.5 spikes/s for either trial type under comparison, were excluded from all analyses.

### Shift function

We used a shift function to quantify if and how two distributions differ (Rousselet et al., 2017). Briefly, using a Harrell-Davis quantile estimator (Harrell and Davis, 1982), distributions were divided into 10 equal parts by 9 “deciles.” For example, the 1^st^ decile is the value below which 10% of the values lie while the 9^th^ decile is the value below which 90% of values lie. The shift function compares a given decile in distribution A with its corresponding decile in distribution B. Corresponding deciles were determined to be significantly different if the confidence interval of their differences, calculated by sampling the difference between bootstrapped distributions 200 times, did not cross 0.

### Activity change during epoch of interest compared to baseline

We calculated the normalized response (*NR*) for each neuron as 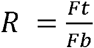, where *Ft* is the median firing rate in the “test” window (either movement preparation epoch or movement initiation epoch) and *Fb* is the median firing rate in the “baseline” window. Since the structure of our task does not include a natural “baseline” epoch – i.e., in which the mouse is in a motionless state unaffected by task demands – our baseline window was defined as the time of odor port entry to reward port exit (i.e., the duration of the whole trial; (Lintz and Felsen, 2016). Neurons with *Fb* < 2.5 spikes/s were excluded from analyses. Statistical significance was determined using a pairwise t-test to compare Ft and Fb from the same trial. Neurons with NR < 1 (p < 0.05) were defined as “Decreasing” and neurons with NR > 1 (p < 0.05) were defined as “Increasing”; all other neurons were categorized as “No Δ” (Fig. 8; Table 1). Note that, by convention, a decreasing neuron that decreases more for contralateral than ipsilateral movement would be considered to have an ipsilateral direction preference (as calculated above), because firing rate is higher for ipsilateral movement (Sato and Hikosaka, 2002; Lintz and Felsen, 2016).

**Figure 7.**
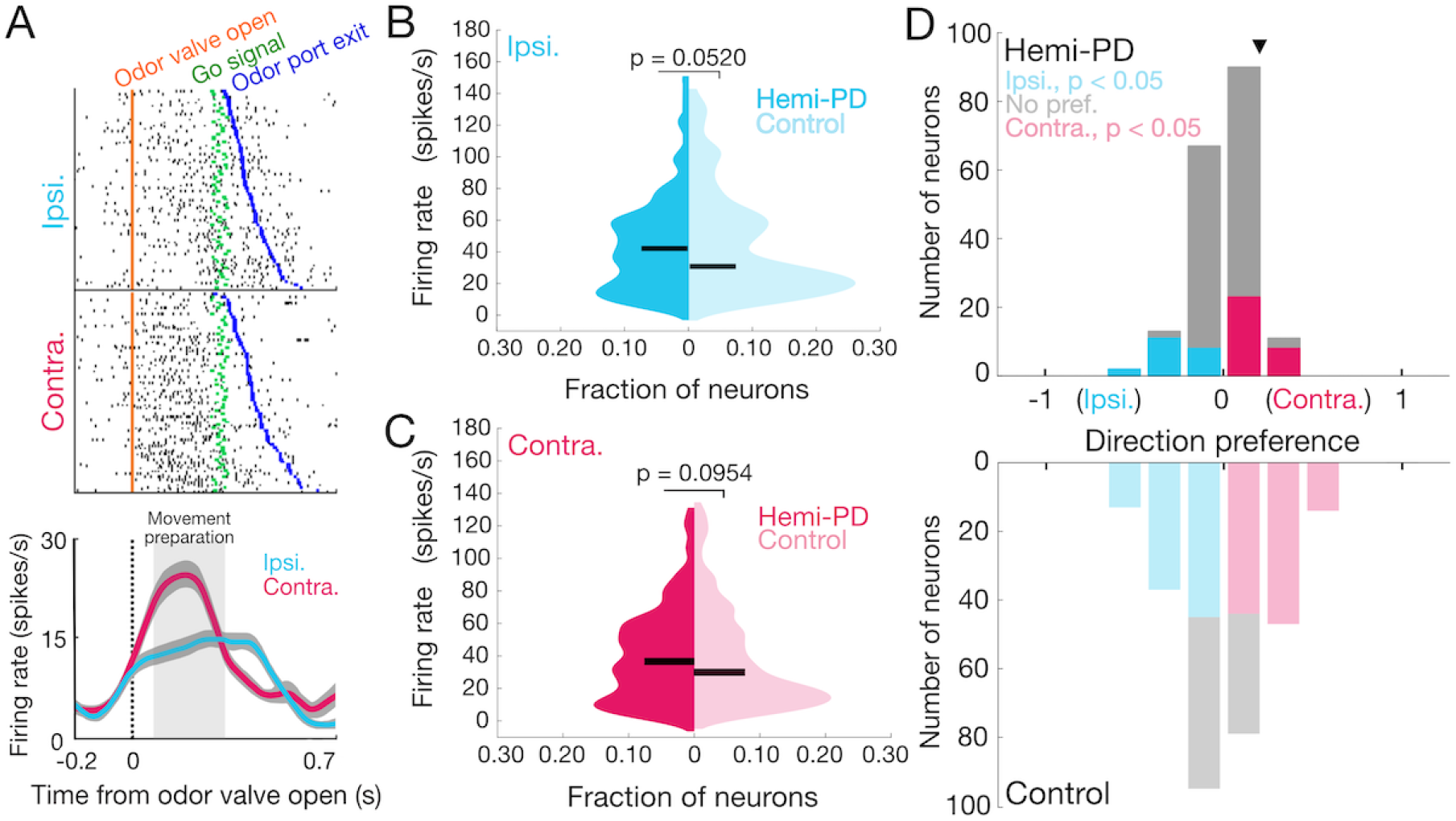
SNr activity during movement preparation in behaving hemi-PD and control mice. ***A***, Rasters (top) and peri-event time histograms (bottom) for an example neuron from a hemi-PD mouse aligned to odor valve open and segregated by choice. Histograms are smoothed with a Gaussian filter; shading, ±SEM. ***B,C*,** Mean firing rate during movement preparation epoch (between odor valve open and go signal) did not differ between populations of neurons in hemi-PD and control mice on ipsilateral (*B*) or contralateral (*C*) trials (p_ipsi_= 0.0520; p_contra_= 0.0954, 1-tailed Wilcoxon rank sum tests; n = 183 neurons in 5 hemi-PD mice; 285 neurons in 4 control mice). Horizontal bars, medians. ***D***, Distribution of direction preferences for population of neurons in hemi-PD mice (top) had a smaller range than in control mice (bottom) (p = 7.31 × 10^-4^, 2-sample Kolmogorov-Smirnov test), and more neurons exhibited a significant preference in control than in hemi-PD mice (p = 2.20 × 10^-16^, χ2-test). Arrowhead, example neuron shown in *A*.

**Figure 8.**
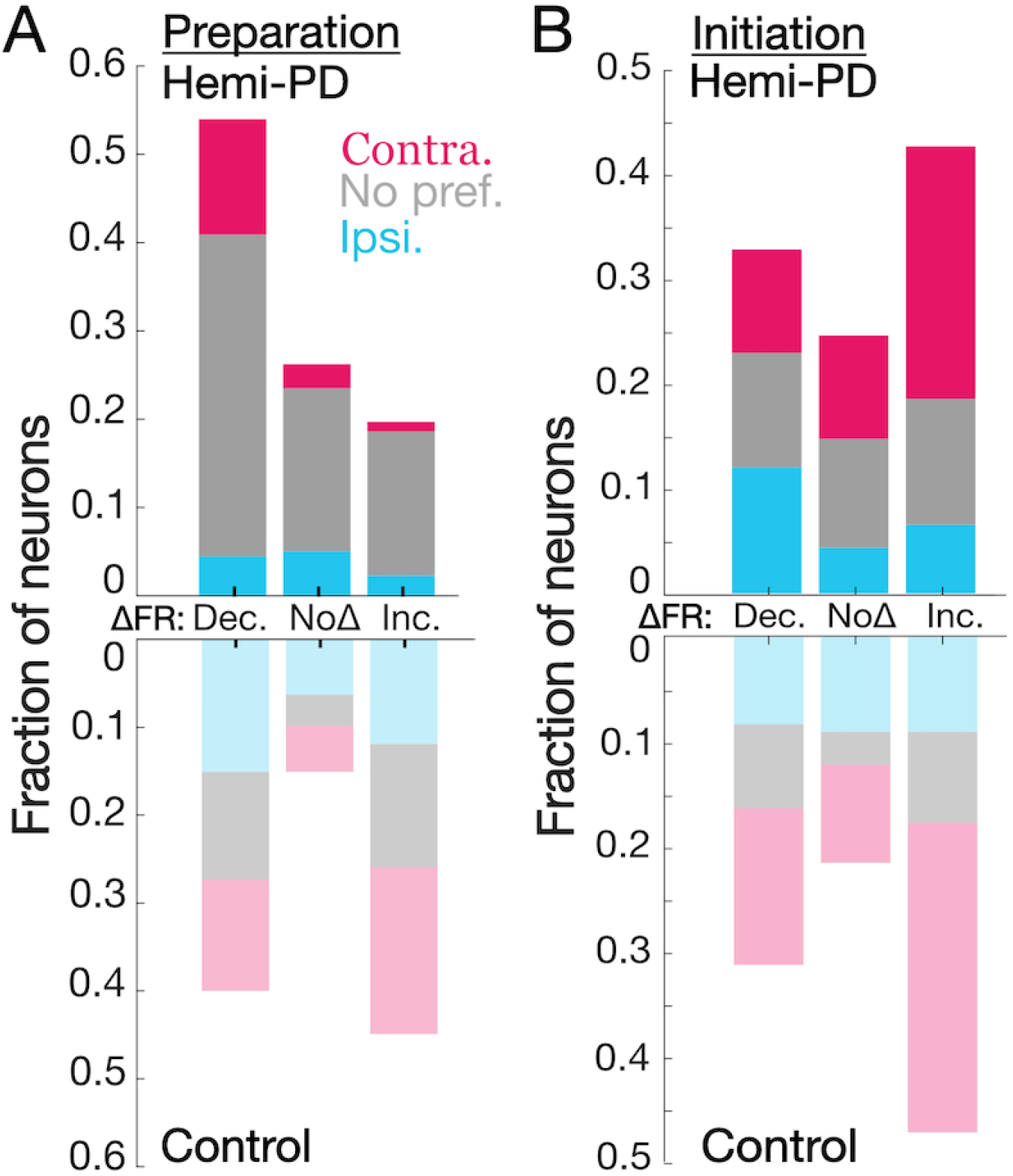
Functional classes of SNr neurons during movement preparation and initiation in behaving hemi-PD and control mice. ***A*,** Fraction of neurons with a given direction preference segregated by whether their average activity during the movement preparation epoch (beginning 100 ms after the odor valve opens and ending with the go signal) significantly increased (Inc.), decreased (Dec.) or did not change (No Δ) relative to average baseline firing rate, for hemi-PD (top) and control (bottom) mice. Proportion of neurons exhibiting an increase, a decrease, and no change differs between hemi-PD and control mice (p = 1.32 × 10^-11^, χ2-test = 50.108, df = 2). Similarly, proportion of ipsilateral, contralateral, and no direction preference neurons differed between hemi-PD and control mice (p = 2.2 × 10^-16^, χ2-test = 149.58, df = 2). ***B*,** Fraction of neurons with a given direction preference segregated by whether their activity during the movement initiation epoch (beginning with the go signal and ending 100 ms after odor poke out) increased, decreased or did not change relative to pre-surgery, for hemi-PD (top) and control (bottom) mice. Proportion of neurons exhibiting an increase, a decrease, and no change did not differ between hemi-PD and control mice (p = 0.459, χ2-test = 1.5586, df = 2; n = 183 neurons in 5 hemi-PD mice; 285 neurons in 4 control mice); proportion of ipsilateral, contralateral, and no direction preference neurons differed between hemi-PD and control mice (p = 4.413 × 10^-5^, χ2-test = 20.049, df = 2).

**Table 1:** Direction preference and activity change from baseline for all SNr neurons recorded from hemi-PD (upper) and control (lower) mice.

| Hemi-PD | Decreased FR | No $\Delta$ FR | Increased FR | Total |
| --- | --- | --- | --- | --- |
| Contralateral | 25 | 5 | 2 | 32 (17.5%) |
| Ipsilateral | 7 | 9 | 4 | 20 (10.9%) |
| Non-selective | 67 | 34 | 30 | 131 (71.6%) |
| Total | 99 (54.1%) | 48 (26.2%) | 36 (19.7%) | 183 neurons |

| Control | Decreased FR | No $\Delta$ FR | Increased FR | Total |
| --- | --- | --- | --- | --- |
| Contralateral | 36 | 15 | 54 | 105 (36.8%) |
| Ipsilateral | 42 | 18 | 34 | 94 (33.0%) |
| Non-selective | 36 | 10 | 40 | 86 (30.2%) |
| Total | 114 (40%) | 43 (15.1%) | 128 (44.9%) | 285 neurons |

### Statistical Analysis

MATLAB was used for all statistical analyses except for *χ*^2^ analyses, which were performed in R. Distributions were tested for normality using the Lilliefors test (lillitest MATLAB function) and unless all data sets under comparison were normally distributed, non-parametric statistical tests were used for that analysis. For consistency, all graphical representations of central tendency are medians, independent of whether parametric or non-parametric statistical tests were used. We used two-tailed tests unless testing for a predicted direction of an effect, in which case one-tailed tests were appropriate.

## Results

### Effect of unilateral dopaminergic cell loss on stimulus-guided and internally-specified movements

To examine how parkinsonian conditions affect stimulus-guided (SG) and internally-specified (IS) movements and their underlying BG output, we first trained mice on a behavioral task designed to elicit these forms of movements (Figs. 1, 2A,B
) (Lintz and Felsen, 2016). Briefly, on each trial of the task, the mouse was presented with a binary mixture of Odors A and B at a central port, waited for an auditory go signal, and moved to the ipsilateral (left) or contralateral (right) reward port for a water reward (Fig. 2A). Each daily experimental session consisted of interleaved blocks of SG and IS trials (Fig. 2B; Materials and Methods) (Lintz and Felsen, 2016). On SG trials, the dominant component of the odor mixture – which varied by trial – determined the side at which reward was delivered: when Odor A was dominant reward was available at the left port, when Odor B was dominant reward was available at the right port, and when Odor A = Odor B reward was equally likely at both ports (Fig. 2B). On IS trials, only the Odor A = Odor B mixture was presented, and reward was delivered at only one port (left or right) throughout the block (Fig. 2B; Materials and Methods). Consistent with previous results (Lintz and Felsen, 2016), the direction of movement on SG trials was selected based on the stimulus (Fig. 2C) while the direction of movement on IS trials was selected based on recent trial history (Fig. 2D).

Upon achieving proficient performance on the task (Fig. 2C,D; Materials and Methods), mice received 6-OHDA injections to one SNc to unilaterally ablate dopaminergic neurons (Ungerstedt, 1976; Chang et al., 2006; Israel and Bergman, 2008; Avila et al., 2010; Smith and Heuer, 2011; Brazhnik et al., 2012; Seeger-Armbruster and von Ameln-Mayerhofer, 2013; Brazhnik et al., 2014) and were implanted with a chronic tetrode drive targeting the ipsilateral SNr to record BG output (Fig. 1). Only mice with > 70% histologically-confirmed dopaminergic cell loss (examined following all behavioral and recording experiments; Fig. 1, Fig. 3A) were included in the “hemi-PD” group for subsequent analyses (5/10 mice; Materials and Methods). Confirming the validity of our hemi-PD model, hemi-PD mice exhibited a greater ipsilateral bias than control mice (saline delivered to SNc) on a standard rotation assay (post-surgery change in net ipsilateral rotations for 5 hemi-PD mice: 16, 53, −6, 151 and 61; for 3 control mice: 6, −11 and −14; p = 0.0357, 1-tailed Wilcoxon rank sum test; Materials and Methods), and exhibited higher mean SNr activity (p = 0.0157, 1-tailed Wilcoxon rank sum test; Fig. 3B), consistent with previous findings in hemi-PD models (Sanderson et al., 1986; Hutchison et al., 1994; Wichmann et al., 1999; Chang et al., 2006; Brazhnik et al., 2014; Lobb and Jaeger, 2015).

Before directly comparing the effects of unilateral dopaminergic cell loss on the SG and IS movements required by our task, we sought to characterize the effects on each trial type individually. Given the rate-model prediction that SNr-recipient nuclei for contralateral movement are over-inhibited (Albin et al., 1989; DeLong, 1990; Obeso et al., 2008; Utter and Basso, 2008; McGregor and Nelson, 2019; Vitek and Johnson, 2019), and supported by our findings of greater ipsilateral bias on the rotation assay and an increase in mean SNr activity (Fig. 3B), we expected hemi-PD mice (as determined by the extent of dopaminergic cell loss; Fig. 3A) to exhibit an ipsilateral bias for both SG and IS movements.

We first examined SG movements. Qualitatively, we found that the psychometric functions fit to post-surgery behavioral sessions (as in Fig. 2) were shifted ipsilaterally relative to pre-surgery sessions in hemi-PD (Fig. 4A), but not control (Fig. 4B), mice. To quantify how bias changed following surgery, we calculated the behavioral directional bias for each post-surgery session and subtracted the pre-surgery bias for that mouse (Materials and Methods). In sessions performed by hemi-PD mice, despite typical levels of variability consistent with other behavioral assays of this 6-OHDA model (Smith and Heuer, 2011), we found that choices after surgery were biased ipsilaterally (p = 8.87 × 10^-8^, 2-tailed Wilcoxon signed rank test) while no post-surgery bias was observed in control mice (p = 0.3480, 2-tailed t-test), and post-surgery bias differed between hemi-PD and control mice (p = 5.64 × 10^-5^, 2-tailed Wilcoxon rank sum test; Fig. 4C). Hemi-PD mice also exhibited changes in reaction time (defined as the time from go signal to reward port entry) consistent with their ipsilateral directional bias. Fig. 4D shows the distribution of reaction times for each trial of all pre-surgery and post-surgery sessions for the same example mouse shown in Fig. 4A. Reaction times are longer post-surgery, most notably for contralateral trials (Fig. 4A, right panel). Control mice also exhibited longer reaction times post-surgery, as expected given the additional weight of the chronic recording drive, but for the example mouse the increase does not appear to be greater for contralateral than ipsilateral movements (Fig. 4E). To quantify the change in reaction time across all mice, we calculated the median reaction time on each post-surgery session and subtracted the overall median pre-surgery reaction time for each mouse (Fig. 4F). As exhibited by the example mice (Fig. 4D,E), reaction times were longer post-surgery, but hemi-PD mice exhibited larger changes in reaction times than control mice (ipsilateral trials: p = 9.90 × 10^-3^, contralateral trials: p = 8.19 × 10^-22^, 2-tailed Wilcoxon rank sum tests; Fig. 4F), and exhibited a larger change on contralateral than on ipsilateral trials (p = 1.66× 10^-12^, 2-tailed Wilcoxon signed rank test). This greater slowing of contralateral movements in hemi-PD mice is consistent with their ipsilateral directional bias (Fig. 4C).

We observed similar effects of unilateral dopaminergic cell loss on IS movements (Fig. 5). For these trials, we first examined bias separately during blocks in which reward was delivered at the ipsilateral port (“ipsilateral blocks”) and at the contralateral port (“contralateral blocks”). Fig. 5A,B shows behavior on IS trials for all pre- and post-surgery sessions for representative hemi-PD and control mice. On ipsilateral blocks, both mice frequently chose the ipsilateral port. On contralateral blocks, however, the hemi-PD mouse was more likely to choose the ipsilateral port post-surgery than pre-surgery (Fig. 5A), while the control mouse did not exhibit this change. Across all mice, we quantified directional bias as we did for SG trials (Materials and Methods), and again found that choices after surgery were biased ipsilaterally in hemi-PD mice (p = 1.49 × 10^-6^, 2-tailed Wilcoxon signed rank test), while no post-surgery bias was observed in control mice (p = 0.3855, 2-tailed t-test); this post-surgery bias differed between hemi-PD and control mice (p = 7.74× 10^-3^, 2-tailed Wilcoxon rank sum test; Fig. 5C). We also found similar effects on reaction times on IS trials as we did on SG trials: hemi-PD mice exhibited larger changes in reaction times than control mice (ipsilateral trials: p = 9.38 × 10^-5^, 2-tailed Wilcoxon rank sum test, contralateral trials: p = 3.90 × 10^-18^, 2-tailed t-test; Fig. 5F), and exhibited a larger change on contralateral than on ipsilateral trials (p = 1.09 × 10^-10^, 2-tailed Wilcoxon signed rank test), again consistent with their ipsilateral directional bias (Fig. 5C). Thus, on both SG and IS trials, unilateral dopaminergic cell loss resulted in fewer and slower contralateral movements, consistent with rate-model predictions.

Next, we sought to examine whether this directional effect was greater in magnitude, indicative of greater impairment, on SG or IS trials. To do so we directly compared, within each session, directional bias on contralaterally-rewarded SG and IS trials. We found that presurgery, hemi-PD mice were more ipsilaterally biased on IS than SG trials (p = 1.99 × 10^-21^, 2tailed Wilcoxon signed rank test; Fig. 6A), as expected since on some SG trials an ambiguous stimulus cue was presented (Fig. 2C). However, after surgery, ipsilateral bias differed little between IS and SG trials (p = 0.0924, 2-tailed Wilcoxon signed rank test; Fig. 6A), resulting in a shift of the distribution of within-session ipsilateral bias differences post-surgery (p = 3.62 × 10^-11^, 2-tailed Wilcoxon rank sum test; Fig. 6B). In contrast, control mice exhibited a greater ipsilateral bias on SG than IS trials both before (p = 1.23 × 10^-8^, 2-tailed Wilcoxon signed rank test) and after (p = 4.72 × 10^-3^, 2-tailed Wilcoxon signed rank test) surgery (Fig. 6C), and the magnitude of this relative bias did not differ between sessions before and after surgery (p = 0.643, 2-tailed Wilcoxon rank sum test; Fig. 6D). Together, these analyses demonstrate that IS trials are relatively more affected than SG trials by unilateral dopaminergic cell loss.

### Effect of unilateral dopaminergic cell loss on basal ganglia output

We sought to determine whether behavioral differences between SG and IS trials could be accounted for by activity in the SNr, which is known to play a role in the movements elicited by this task (Lintz and Felsen, 2016). In the same mice described in the above behavioral results, we used chronically implanted tetrodes to record from 183 SNr neurons in hemi-PD mice (n = 5) and 285 neurons in control mice (n = 4) that met our analysis criteria (Materials and Methods). We focused on activity underlying movement preparation, which has been implicated in parkinsonian motor deficits (Dick et al., 1984; Jahanshahi et al., 1992; Suri et al., 1998; Berardelli et al., 2001; Cutsuridis and Perantonis, 2006; Moroney et al., 2008; Wu et al., 2015; Hess and Hallett, 2017). Consistent with previous results (Lintz and Felsen, 2016), we observed that SNr activity recorded from hemi-PD mice was often modulated during the “movement preparation epoch” (from 100 ms after the odor valve was opened until the go signal) and depended on movement direction (Fig. 7A). We first asked whether firing rate during this epoch differed between hemi-PD and control mice, similar to the difference we and others have observed in baseline activity (Fig. 3B) (Hutchison et al., 1994; Wichmann et al., 1999; Galati et al., 2010; Wang et al., 2010a; Seeger-Armbruster and von Ameln-Mayerhofer, 2013; Brazhnik et al., 2014; Lobb and Jaeger, 2015; Aristieta et al., 2016; Willard et al., 2019). The rate model would predict elevated SNr activity in hemi-PD mice, consistent with the ipsilateral bias that we observed given that SNr activity inhibits downstream motor centers that primarily mediate contralateral movement. However, we found no overall difference between groups for movement in either direction (p_ipsi_= 0.0520; p_contra_= 0.0954, 1-tailed Wilcoxon rank sum tests; Fig. 7B,C). The ipsilateral bias exhibited by hemi-PD mice on SG and IS trials therefore cannot be explained by an *absolute* increase in SNr activity during movement preparation, as would be predicted by the simplest form of the rate model.

We next examined whether unilateral dopaminergic cell loss affected the *relative* activity of individual neurons between ipsilateral and contralateral movements (Fig. 7A), which could also potentially account for the ipsilateral behavioral bias. We therefore calculated the “direction preference” of each neuron, which quantifies the difference in firing rate during a specified epoch between ipsilateral and contralateral movements, and ranges from −1 (higher firing rates for ipsilateral movements) to 1 (higher firing rates for contralateral movements), where 0 represents no preference (Materials and Methods) (Green and Swets, 1966; Feierstein et al., 2006; Lintz and Felsen, 2016). We found that the populations of neurons recorded in hemi-PD and control mice each exhibited a range of preferences, with some neurons preferring ipsilateral movement (Fig. 7D, cyan; p < 0.05, permutation test; Materials and Methods) and some preferring contralateral movement (Fig. 7D, magenta; p < 0.05, permutation test). However, the distributions of preferences exhibited by neurons recorded in hemi-PD and control mice differ (p = 7.31 × 10^-4^, 2-sample Kolmogorov-Smirnov test) in two key respects.

First, the population of SNr neurons in hemi-PD mice exhibited weaker direction preference than the population in control mice. We quantified this difference with several complementary analyses. A smaller proportion of neurons in hemi-PD mice (52/183, 28%; Table 1) than control mice (199/285, 70%; Table 1) exhibited a significant direction preference (p = 2.20 × 10^-16^, χ2-test = 147, df = 1). When the entire population in hemi-PD and control mice is considered (i.e., including neurons with and without a significant direction preference), the strength of the preference, independent of sign, was smaller in hemi-PD mice (median = 0.0815) than control mice (median = 0.179; p = 1.87× 10^-12^, 1-tailed Wilcoxon rank sum tests). Finally, we used a shift function analysis to identify the deciles in which the distributions differed (Materials and Methods) (Rousselet et al., 2017). This analysis revealed that preferences were closer to 0 in hemi-PD than control mice in the 3 deciles representing the most ipsilateral preferences and in the 3 deciles representing the most contralateral preferences, consistent with weaker direction preference in hemi-PD mice. Together, these analyses indicate that unilateral dopaminergic cell loss results in a fundamental disruption of the representation of movement direction that is normally observed in the SNr (Handel and Glimcher, 1999; Berardelli et al., 2001; Lintz and Felsen, 2016).

Second, the population of SNr neurons in hemi-PD mice, but not control mice, exhibited a slight bias towards contralateral preferences. While the entire distribution of preferences in hemi-PD mice was not contralaterally skewed (median = 0.136; p = 0.368, 1-tailed Wilcoxon signed rank test), of the neurons that exhibited a significant preference (Fig. 7D, magenta and cyan, Table 1), more were contralateral-preferring (32/52, magenta; Table 1) than ipsilateral-preferring (20/52, cyan; p = 0.0155, χ2-test = 4.65 df = 1; Table 1). In contrast, in control mice we found roughly equal proportions of contralateral-preferring (105/199, magenta; Table 1) and ipsilateral-preferring (94/199, cyan; Table 1) neurons (p = 0.158, χ2-test = 1.01, df = 1; Fig. 7D, Table 1). This analysis indicates that, in the population of SNr neurons in which direction preference is spared, unilateral dopaminergic cell loss results in more neurons exhibiting higher activity for contralateral than ipsilateral movements, consistent with the ipsilateral behavioral bias that we observed (Figs. 4, 5, Table 1).

To gain insight into the reorganization of BG output induced by unilateral dopaminergic cell loss, we next examined these systematic differences in preferences between hemi-PD and control mice within functional classes of SNr neurons (Fig. 8, Table 1). Separate subpopulations of SNr neurons are known to exhibit increases or decreases in activity as movements are prepared and initiated (Handel and Glimcher, 1999; Sato and Hikosaka, 2002; Lintz and Felsen, 2016); these subpopulations presumably play different functional roles. We therefore categorized neurons into one of three classes based on whether their activity during movement preparation increased, decreased, or did not change, relative to baseline (Table 1; Materials and Methods). We first noticed a clear difference in the proportion of neurons in each class between hemi-PD and control groups (p = 1.32 × 10^-11^, χ2-test = 50.108, df = 2; n = 183 neurons, 5 hemi-PD mice; n = 285 neurons, 4 control mice; Fig. 8A, Table 1). Specifically, we found that a higher proportion of neurons recorded in hemi-PD mice (54.1%) than in control mice (40.0%) exhibited decreased activity, and a corresponding lower proportion of neurons recorded in hemi-PD mice (19.7%) than in control mice (44.9%) exhibited increased activity (Fig. 8A, Table 1; Materials and Methods). In addition, our finding that fewer neurons in hemi-PD than control mice exhibited a significant direction preference held true across all 3 subpopulations of neurons (Fig. 8A, Table 1). Finally, we found that the contralateral bias in the hemi-PD group among neurons exhibiting a direction preference was largely due to those that exhibited decreased activity during movement preparation (contralateral:ipsilateral ratio = 25:7, p = 1.07 × 10^-5^, χ2-test = 18.1, df = 1; Fig. 8A, Table 1); this subpopulation in the control group did not show this effect (contralateral:ipsilateral ratio = 36:42; p = 0.788, χ2-test = 0.641, df = 1; Fig. 8A, Table 1). Thus, unilateral dopaminergic cell loss resulted in a larger proportion of SNr neurons that release downstream motor centers from inhibition, with a greater effect on ipsilateral than contralateral movements, accounting for the relationship between SNr activity and the ipsilateral behavioral bias.

Thus far we have focused on SNr activity during the movement preparation epoch. As our task is designed to separate movement preparation from initiation, we were able to extend our analyses to movement initiation. We expected to observe similar changes during movement initiation, consistent with previous studies (Kravitz et al., 2010; Wang et al., 2010b; Abedi et al., 2013; Freeze et al., 2013). We therefore repeated our firing rate, direction preference and functional class analyses for the period from the go signal until 100 ms after odor port exit, when the movement is initiated. As with the movement preparation epoch, we found no overall difference in firing rate during movement initiation between the hemi-PD and control groups for movement in either direction (p_ipsi_= 0.193; p_contra_ 0.103, 1-tailed Wilcoxon rank sum tests). Next, as with movement preparation, we found that a smaller proportion of neurons in hemi-PD than control mice exhibited a direction preference during movement initiation (p = 4.41 × 10^-5^, χ2-test = 20.0, df = 2; Fig. 8B), but our shift function analysis comparing the distributions of direction preferences in hemi-PD and control mice revealed only one differing decile (8^th^ decile). Consistent with this result, we found no difference between hemi-PD and control mice in the proportion of neurons that increased or decreased activity during movement initiation compared to baseline (p = 0.459, χ2-test = 1.56, df = 2; n = 183 neurons, 5 hemi-PD mice; n = 285 neurons, 4 control mice; Fig. 8B, Table 1). Thus, dopaminergic cell loss appears to affect BG output more during movement preparation than during movement initiation in the context of our behavioral task.

Finally, we examined whether the changes in SNr activity during movement preparation associated with unilateral dopaminergic cell loss were consistent with the stronger ipsilateral behavioral bias on IS compared to SG trials (Fig. 6). For example, the representative neuron shown in Fig. 9A appears to exhibit a contralateral preference, consistent with an ipsilateral behavioral bias, on IS but not SG trials. To examine this phenomenon across the population, we compared direction preference during the movement preparation epoch between IS and SG trials within the same session (Fig. 9B). In the hemi-PD group, we found that preference was significantly greater (i.e., more contralateral) on IS than SG trials (p = 0.00390, 1-tailed Wilcoxon signed rank test, n = 158 neurons; Fig. 9B). We also observed a significant difference in control mice (p = 0.0180, 2-tailed Wilcoxon signed rank test; n = 285 neurons), but in the opposite direction (i.e., more ipsilateral). Together, these analyses of SNr activity show that changes in BG output in hemi-PD mice are consistent with their overall ipsilateral behavioral bias (Figs. 4, 5), as well as their stronger bias on IS than SG trials (Fig. 6).

**Figure 9.**
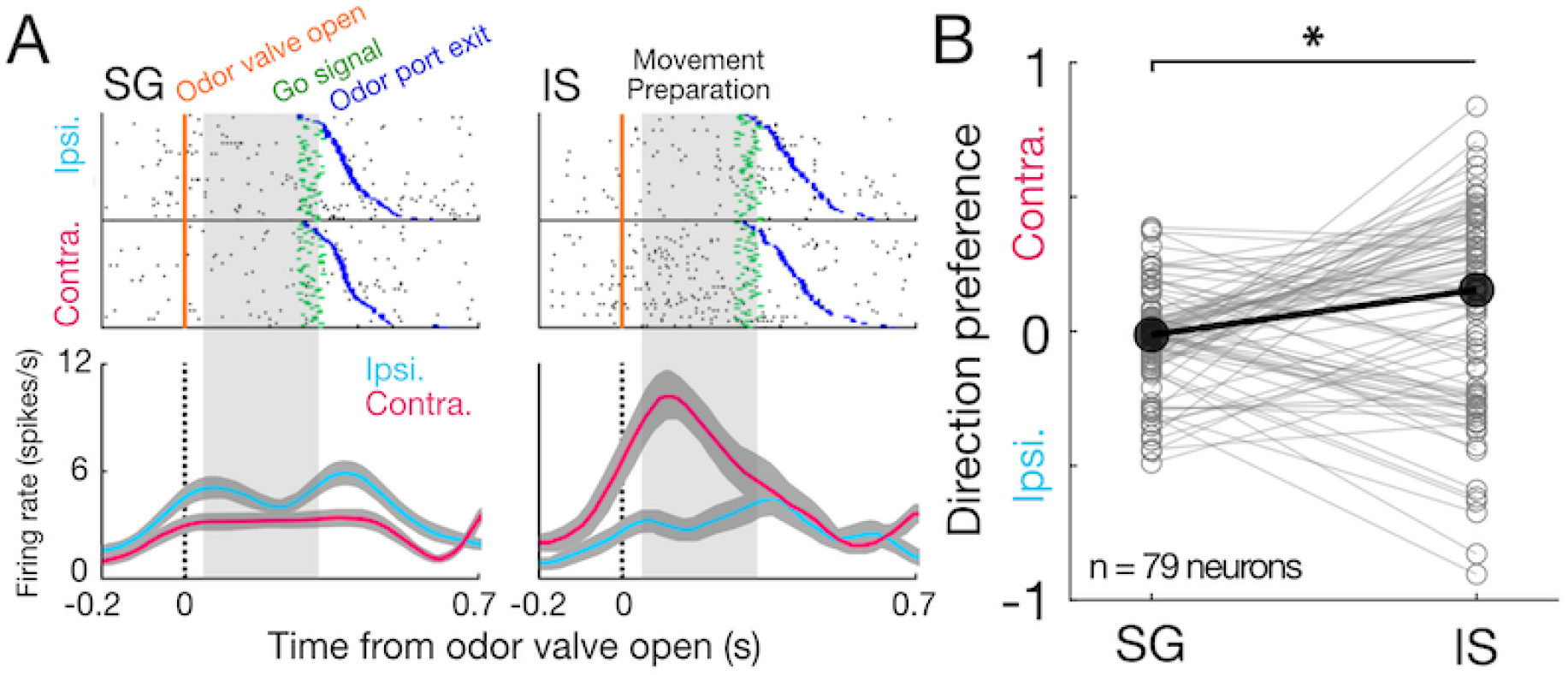
SNr activity on stimulus-guided (SG) and internally-specified (IS) trials in hemi-PD mice. ***A*,** Rasters (top) and peri-event histograms (bottom), on stimulus-guided (SG, left) and internally-specified (IS, right) trials, for an example neuron from a hemi-PD mouse aligned to odor valve open and segregated by choice. Histograms are smoothed with a Gaussian filter; shading, ±SEM. ***B*,** Direction preference on SG and IS trials for population of SNr neurons in hemi-PD mice. Each neuron is represented by a pair of connected gray symbols; only neurons with significant preference (p < 0.05) on SG and/or IS trials are shown. Preference was more contralateral on IS than SG trials (p = 0.0132, 1-tailed Wilcoxon signed rank test, n = 79). Black symbols, medians.

## Discussion

In this study, we primarily sought to determine whether BG output could explain the greater impairment in IS than SG movements under parkinsonian conditions. By recording SNr activity in hemi-PD mice performing SG and IS movements, we found that unilateral dopaminergic cell loss alters the relationship between SNr activity and movement direction as movements are prepared (Figs. 7D, 8, Table 1), consistent with the ipsilateral bias in behavior (Figs 4, 5). While we did not observe an absolute increase in preparation-related SNr activity in hemi-PD mice (Fig. 7B,C), as predicted by the classical rate model of parkinsonian BG activity (DeLong, 1990; McGregor and Nelson, 2019), our findings are consistent with a direction-sensitive form of the rate model in which greater SNr output inhibits downstream motor nuclei mediating contralateral movements (DeLong, 1990; McGregor and Nelson, 2019; Vitek and Johnson, 2019). In contrast, neural activity during movement initiation was little changed between hemi-PD and control groups (Fig. 8B), suggesting that parkinsonian conditions affect BG output subserving movement preparation more than initiation, consistent with studies in PD patients (Jahanshahi et al., 1992; Beiser and Houk, 1998; Suri et al., 1998; Cutsuridis and Perantonis, 2006; Moroney et al., 2008; Wu et al., 2015).

While unilateral dopaminergic cell loss resulted in an ipsilateral bias on both SG and IS trials (Figs. 4 and 5), we found that the effect was larger on IS trials (Fig. 6). This difference was reflected in the activity of SNr neurons, which exhibited a stronger contralateral preference (i.e., higher activity on contralateral than ipsilateral trials) on IS than SG trials (Fig. 9). This finding is consistent with our previous work showing that SNr activity is more sensitive to movement direction under IS than SG conditions (Lintz and Felsen, 2016), which suggests that the cognitive processes associated with IS movements are more BG-dependent than those associated with SG movements. One key factor that may explain the difference in BG-dependence between IS and SG trials is the influence of prior choices and outcomes, which are crucial for determining the correct direction of movement on IS, but not SG, trials. The SNr receives direct excitatory input from the pedunculopontine tegmental nucleus (Scarnati et al., 1984; Beninato and Spencer, 1987), which encodes prior choices and outcomes (Thompson and Felsen, 2013); SNr activity itself reflects prior choices (Lintz and Felsen, 2016), and in general the BG are thought to bias activity in downstream motor centers towards movements associated with larger rewards (Hikosaka et al., 2000; Sato and Hikosaka, 2002; Watanabe et al., 2003; Kawagoe et al., 2004; Hikosaka et al., 2006). The disruption of BG signaling by unilateral dopaminergic cell loss may therefore affect the (adaptive) influence of priors on movement selection (Perugini et al., 2016), resulting in a greater deficit on IS than SG trials. The importance of the BG in representing priors is also consistent with our finding that SNr activity related to movement preparation, which is influenced by priors, was more affected than activity related to movement initiation, which is not.

In addition to the shift in the population of neurons in hemi-PD mice towards contralateral preference consistent with the ipsilateral behavioral bias, a large proportion of neurons exhibited no relationship between activity and movement direction (gray bars in Figs. 7D, 8A, Table 1). Further, we found that more neurons in hemi-PD mice exhibited a decrease than an increase in activity during movement preparation compared to baseline activity, which was not the case in control mice (Fig. 8A, Table 1), perhaps due to the increased baseline activity in hemi-PD mice (Fig. 3B). While these finding do not directly relate to our primary behavioral readout of directional bias, they suggest a profound reorganization of BG output under parkinsonian conditions. How this reorganization could contribute to other features of parkinsonian behavior can be examined in future studies.

While we have demonstrated a link between the electrophysiological and behavioral effects of unilateral dopaminergic cell loss, it is worth considering potential caveats in interpreting our results. First, it is possible that compensatory mechanisms unrelated to the electrophysiological effects of unilateral dopaminergic cell loss are responsible for the behavioral bias we observed, particularly since the hemi-PD mice included in our study all exhibited a behavioral bias in the same direction (ipsilateral). While our electrophysiological results are consistent with this bias, we would be able to draw stronger conclusions about the relationship between BG output and behavior with a set of mice exhibiting a wider range of behavioral biases. Indeed, one mouse that was excluded from our hemi-PD group due to insufficient dopaminergic cell loss (< 70%) exhibited a significant *contralateral* bias on both SG (p = 3.10 × 10^-3^, 2-tailed Wilcoxon signed rank test, median bias = −0.223) and IS trials (p = 5.41 × 10^-4^, 2-tailed Wilcoxon signed rank test, median bias = −0.0798). When we analyzed the SNr recordings from this mouse, we found that the neurons with a significant direction preference (74/83 neurons) largely exhibited an *ipsilateral* preference during the movement preparation epoch (ipsilateral:contralateral ratio = 59:15, p = 7.74 × 10^-13^, χ2-test = 50.0, df = 1, see Fig. 8A, Table 1 for comparisons). While we can only speculate about the potential compensatory adaptations that emerge in the BG with relatively moderate dopaminergic cell loss, and we cannot rule out contributions of additional mechanisms to the behavioral bias, the fact that the direction of behavioral bias remains consistent with neural direction preference further supports our finding that BG output mediates the effect of parkinsonian conditions on behavior.

Second, olfactory deficits are a known hallmark of PD (Hawkes and Shephard, 1993; Hawkes, 1995; Tarakad and Jankovic, 2017; Tekriwal et al., 2017). Given our use of an olfactory task, is it possible that the behavioral effects under parkinsonian conditions are due to sensory and not motor deficits. We suggest that this is unlikely for two reasons: 1) on IS trials the olfactory cue was not informative about reward location, 2) and on SG trials a deficit in olfactory discrimination would be reflected in a flattening, rather than a shift, in the psychometric function, which we did not observe.

Finally, while the hemi-PD model provides a powerful approach for examining the neural basis of parkinsonian movement (Ungerstedt, 1976; Avila et al., 2010; Galati et al., 2010; Brazhnik et al., 2012; Brazhnik et al., 2014; Dorval and Grill, 2014), PD typically presents with bilateral dopaminergic cell loss. It is possible that dopaminergic input from the spared SNc may compensate for the unilateral insult, and other differences between the model and the clinical condition must be considered. However, our comparison between ipsilateral and contralateral movements, at the behavioral and neurophysiological levels, was well suited to the hemi-PD model, and we suggest that our results can cautiously inform the reorganization of BG output that occurs in PD.

In conclusion, we found that the behavioral effects of unilateral dopaminergic cell loss, including differences between stimulus-guided and internally-specified movements, can be accounted for by changes in SNr activity during movement preparation. While our results could not be explained by the simplest rate-model prediction that BG output is tonically elevated by dopaminergic cell loss, they were nevertheless consistent with a form of the rate model in which movement direction is influenced by the rate of BG output during movement preparation. Future studies can examine how the changes in SNr activity observed here affect the activity of SNr-recipient structures, as well as how the BG interact with other motor systems to differentially mediate stimulus-guided and internally-specified movements under parkinsonian conditions.

## Acknowledgments

Technical support was provided by the Optogenetics and Neural Engineering Core at the University of Colorado School of Medicine, funded in part by the National Institutes of Health (P30NS048154). We thank Quang Dang and Ben Peterson for data collection, and members of the Felsen lab for comments on the manuscript.

## Funding

This work was supported by the National Institutes of Health (R01NS079518, P30NS048154); and the Boettcher Foundation (Webb-Waring Biomedical Research Award).

## Declarations of interest

none

## Notes

### Competing Interest Statement

The authors have declared no competing interest.

